# An unexpected histone chaperone function for the MIER1 histone deacetylase complex

**DOI:** 10.1101/2022.07.22.501112

**Authors:** Siyu Wang, Louise Fairall, Khoa Pham, Timothy J Ragan, Dipti Vashi, Mark O. Collins, Cyril Dominguez, John W.R. Schwabe

**Affiliations:** Institute for Structural and Chemical Biology, University of Leicester, Leicester. LE1 7RH. UK; School of Biosciences, University of Sheffield, Sheffield. S10 2TN. UK; biOMICS facility, Mass Spectrometry Centre, University of Sheffield, Sheffield. S10 2TN. UK

## Abstract

Histone deacetylases 1 and 2 (HDAC1/2) serve as the catalytic subunit of six distinct families of nuclear complexes. These complexes repress gene transcription through removing acetyl groups from lysine residues in histone tails. In addition to the deacetylase subunit, these complexes typically contain transcription factor and/or chromatin binding activities. The MIER:HDAC complex has hitherto been poorly characterized. Here we show that MIER1 unexpectedly co-purifies with an H2A:H2B histone dimer. We show that MIER1 is also able to bind a complete histone octamer. Intriguingly, we found that a larger MIER1:HDAC1:BAHD1:C1QBP complex additionally co-purifies with an intact nucleosome on which H3K27 is either di- or tri-methylated. Together this suggests that the MIER1 complex acts downstream of PRC2 to expand regions of repressed chromatin and to deposit histone octamer onto nucleosome-depleted regions of DNA.

## Introduction

The histone deacetylase enzymes, HDACs 1 and 2 act as epigenetic “erasers” removing acetyl groups from histone tails (*1*). This results in the loss of binding sites for regulatory “reader” proteins and causes chromatin compaction by restoring the positive charge on histone tails. HDACs 1 and 2 are highly homologous enzymes that are assembled into six distinct multiprotein complexes: NuRD, CoREST, Sin3A, MiDAC, RERE and MIER (*2-7*). There appears to be little or no redundancy between these complexes since mice lacking unique components typically die during embryogenesis (*5, 8-10*). Apart from the HDAC enzyme, the complexes differ in subunit composition, but often contain chromatin and/or transcription factor interaction activities. The MIER complex has to date received relatively little attention, but is likely to have an important regulatory function since it is present from nematodes to man.

The MIER complex is named after the MIER co-repressor scaffold proteins that directly recruit HDAC1/2 through a canonical ELM2-SANT domain (*11*). There are three paralogous genes *mier*1, 2 & 3. MIER1, also known as Mesoderm Induction Early Response protein 1, was first isolated as a fibroblast growth factor regulated gene in *Xenopus laevis* (*4, 12*). The MIER proteins are predominantly located in the nucleus (*12*). The central ELM2-SANT domain of MIER1, 2 and 3 is highly conserved; the amino-terminal region is moderately conserved and the carboxy-terminal region is poorly conserved.

Several studies have shown that the BAH domain-containing protein, BAHD1, interacts with the MIER1:HDAC1 complex (*13, 14*). BAH domains are found in many chromatin-associated proteins including several proteins involved in transcriptional repression such as SIR3, MTA1, RERE, BAHCC1 and BAHD1 (*15, 16*). BAH domains have been shown to mediate interaction both with intact nucleosomes and with specific histone marks such as H4K20me2 and H3K27me3 (*17-20*). Both BAHCC1 and BAHD1 recognise H3K27me3 suggesting that they are “reader” domains acting downstream of the PRC2 complex (*10, 21*). It is unclear, however, whether these BAH domains simply recognise the H3K27me3 histone mark, or whether they can bind to intact nucleosomes in an analogous fashion to the BAH-containing proteins ORC1 and SIR3 (*17, 20*).

Here we report a biochemical and functional study of the MIER1:HDAC1 and MIER1:HDAC1:BAHD1 complexes. We find that the MIER1 amino-terminal domain mediates interaction with histones H2A and H2B and is able to bind to an intact histone octamer. The MIER1:HDAC1:BAHD1 complex co-purifies with the multi-functional protein C1QBP and with an intact nucleosome specifically bearing H3K27me2/3 marks. The finding that a histone deacetylase complex acts as a histone chaperone is unexpected, but makes sense in terms of increasing nucleosome density as part of the transcriptional repression activity.

## Results

### MIER1 binds to HDAC1/2 through the ELM2-SANT domain

We have previously shown that the ELM2-SANT domains from MTA1, MIDEAS and RCoR1 mediate interaction of the NuRD, MiDAC and CoREST complexes with HDACs 1 and 2 (*5-7*). To confirm that the homologous ELM2-SANT domain from MIER1 is also capable of binding HDAC1/2 we co-expressed FLAG-tagged MIER1(aa:171-350) with full-length HDAC1 in HEK293F cells (Fig. 1). Affinity purification on anti-FLAG resin, followed by size exclusion chromatography confirmed the expected interaction (Fig. 1b). The elution profile is consistent with a monomeric 1:1 complex. This was confirmed using size exclusion chromatography coupled with multi-angle light scattering (SEC-MALS). The complex molecular weight was determined to be 73.6±1.5 kDa (Fig. 1c), which is consistent with a monomeric complex (HDAC1 55kDa + MIER1(aa:171-350) 21kDa) and contrasts with the dimeric MTA1 and tetrameric MIDEAS complexes (*5, 7*). HDAC assays confirmed that the complex is active; can be further activated by inositol-hexakis-phosphate and is inhibited by hydroxamic and benzamide-based inhibitors (Fig. 1d).

**Fig. 1.**
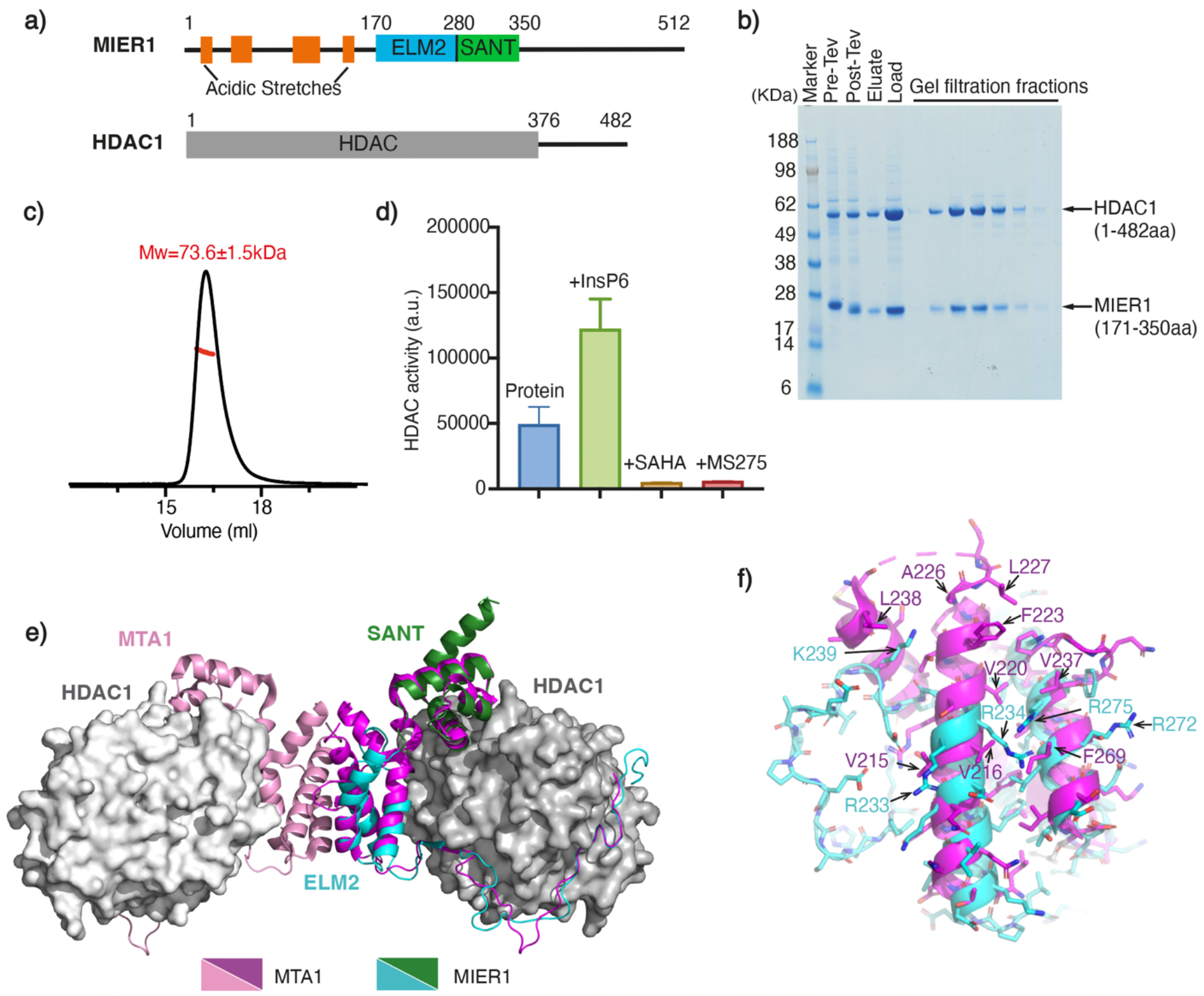
The ELM2-SANT domain of MIER1 co-purifies with HDAC1 and forms a stable, stoichiometric, enzymatically active complex. a) Schematic of the domain structure of full-length MIER1 and HDAC1. b) Purification of the FLAG-tagged MIER ELM2-SANT (aa:171-350):HDAC1 complex using Anti-FLAG resin followed by a Superdex S200 size exclusion column shown on a SDS-PAGE gel. c) Determination of the stoichiometry/molecular weight of the MIER1 ELM2-SANT (aa:171-350):HDAC1 complex using SEC-MALS. The theoretical molecular weight is 75 kDa and the measured molecular weight for 1:1 stoichiometry is 73.6±1.5 kDa. d) Deacetylase activity of the MIER1 ELM2-SANT (aa:171-350):HDAC1 complex in the presence of 100 μM Ins(1,2,3,4,5,6)P_6_ (InsP_6_), 30 μM SAHA and 30 μM MS275. The background of the assay with no complex has been subtracted. Error bars indicate the SEM (n = 3). e) Alphafold2 multimer prediction for the ELM2-SANT domain of MIER1(aa:171-350) (cyan-forest green) with HDAC1(grey) complex compared with the crystal structure of the MTA1(aa:162-354) (pink/magenta):HDAC1(grey) complex (*46*). f) Dimerisation interface of ELM2 domain from MTA1(magenta) compared with ELM2 domain of MIER1(cyan).

We used Alphafold2 multimer (*22*) to predict the structure of the MIER1-ELM2-SANT domain bound to HDAC1 (Fig. 1e). The region of the ELM2 domain that mediates dimerization in MTA1 is clearly unable to support dimerization in MIER1. Helix 2 is absent in MIER1 and the non-polar residues located at the MTA1 dimer interface are mostly replaced with basic residues in MIER1(Fig. 1f).

### The amino terminal region of MIER1 co-purifies with endogenous H2A:H2B histone dimers

To characterise the native complex, we co-expressed FLAG-tagged full-length MIER1, together with full-length HDAC1. Following purification, we observed the expected 1:1 interaction with HDAC1 (Fig. 2a). Unexpectedly however, in addition to the bands for MIER1 and HDAC1, we observed two low molecular-weight proteins that co-purify with the complex through both the affinity and size exclusion purifications. Mass-spectrometry (MS) analysis of these bands identified these proteins to be the endogenous histones H2A and H2B (Fig. 2a).

**Fig. 2.**
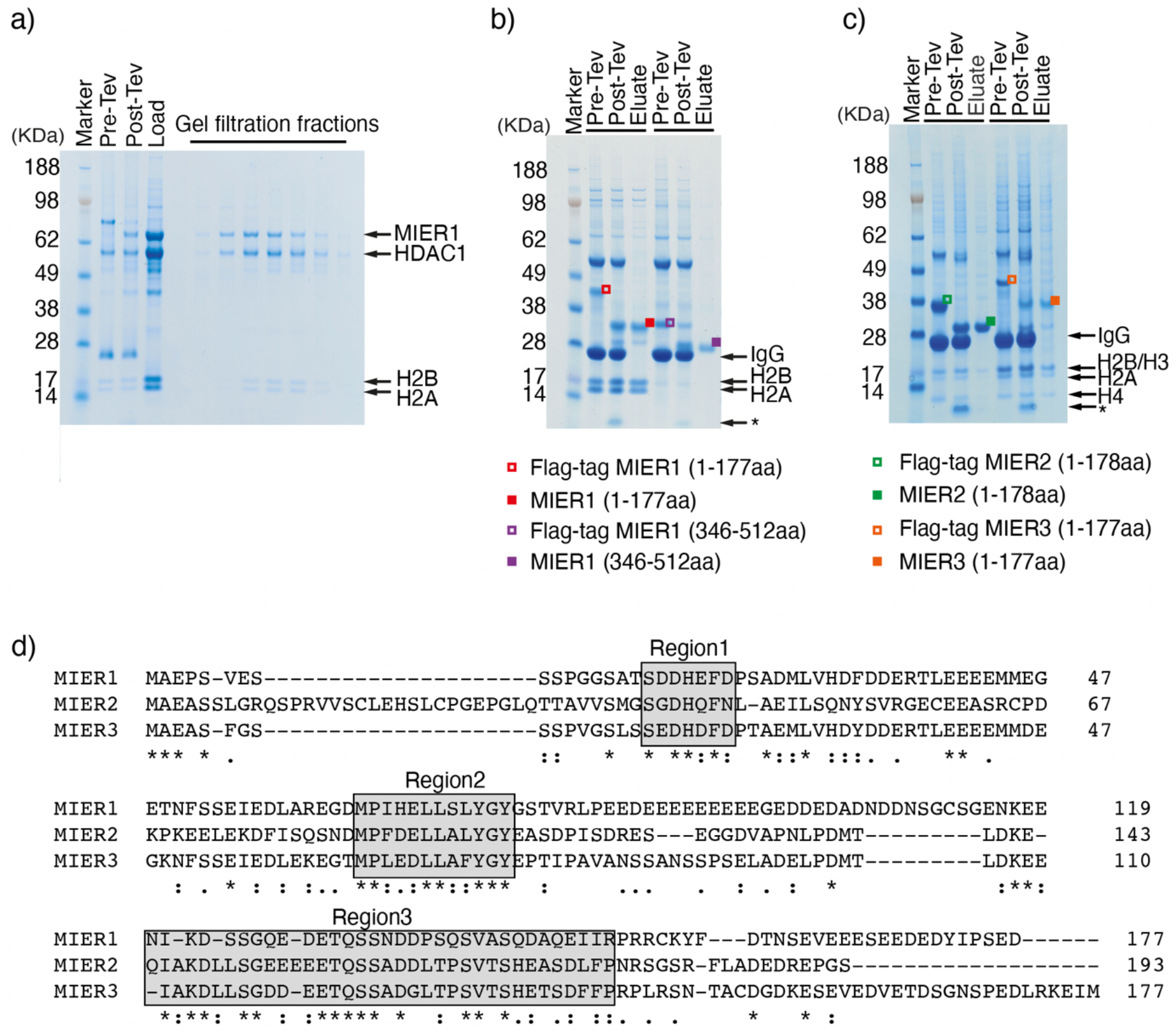
Interaction of MIER 1, 2 and 3 with a histone H2A:H2B dimer. a) Full length FLAG tagged MIER1(aa:1-512):HDAC1 complex co-purifies with endogenous H2A and H2B. SDS-PAGE showing the purification with Anti-FLAG resin and the fractions from the Superdex S200 size exclusion column. b) The amino-terminus of MIER1(aa:1-177) (LHS red symbols) interacts with a histone H2A:H2B dimer. The C-terminus of MIER1(aa:346-512) (RHS purple symbols) shows no interaction. c) MIER2(aa:1-178) (LHS green symbols) and MIER2(aa:1-177) (RHS orange symbols) co-purify with endogenous H2A:H2B. d) Sequence alignment of amino terminus of MIER1, MIER2 and MIER3. Identical and conserved residues are highlighted in grey boxes. Identical residues are labelled with stars, strong similarity is labelled with two dots and a single dot indicates weak similarity.

This unexpected interaction with histones H2A and H2B was not observed with the isolated ELM2-SANT domain (Fig. 1b) suggesting that either the amino-terminal or carboxy-terminal regions mediate this interaction. To identify the interaction region, we expressed FLAG-tagged MIER1(aa:1-177) and FLAG-tagged MIER1(aa:346-512). The H2A:H2B dimer co-purified with the amino-terminal construct and not with the carboxy-terminal construct. Furthermore, the coomassie staining is consistent with a stoichiometric 1:1:1 complex (Fig. 2b).

MIER1 has two paralogues MIER2 and MIER3. Whilst the ELM2-SANT domain is highly conserved (63% identity), the amino-terminal region is less well conserved (Fig. 2d). To determine whether histone binding is a conserved feature of all three proteins, we expressed the amino-terminal domains of MIER2 and MIER3 in HEK293 cells. Both MIER2 and MIER3 co-purified with histones H2A:H2B (Fig. 2c) suggesting histone binding is a common and important property of this family of HDAC co-repressor proteins. Interestingly, in this experiment, histones H3 and H4 also co-purify with the MIER proteins (see below).

We infer from the fact that the H2A:H2B heterodimer co-purifies with MIER1 over both affinity and size-exclusion columns that the interaction must be relatively strong, or at least the off-rate is slow. Using isothermal calorimetry we measured a dissociation constant of approximately 400 nM (Fig. 3a).

**Fig. 3.**
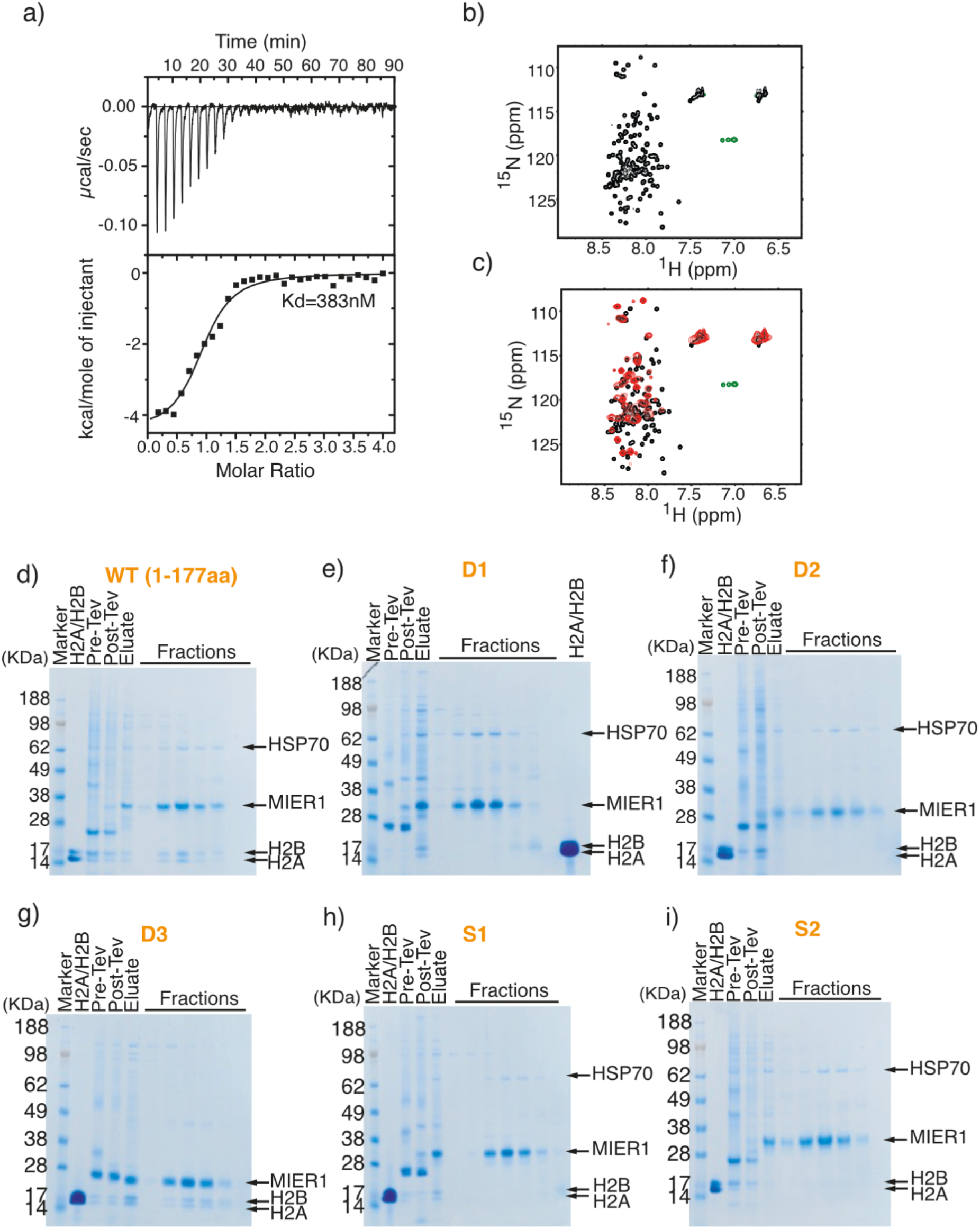
Characterisation of the interaction between amino-terminus of MIER1(aa:1-177) and endogenous H2A:H2B dimer. a) Isothermal titration calorimetry of MIER1(aa:1-177) binding to histones H2A:H2B Raw calorimetric outputs are shown on top and binding isotherms describing formation of the MIER1(aa:1-177):H2A:H2B complex are shown at the bottom. b) (^1^H-^15^N) HSQC experiment of the labelled MIER1(aa:1-177) protein in 1x PBS pH7.4 and 0.5 mM TCEP. c) HSQC results of MIER1(aa:1-177) with the addition of histone H2A and H2B (red) at ratio 1:1 compared with free MIER1(black). d-i) The Flag-tagged wild type MIER1(aa:1-177) and scrambled/deletion regions of MIER1 were expressed in the HEK 293F cells, purified on Anti-FLAG resin followed by a Superdex S200 column. D1, D2, D3, S1 or S2 are the deleted or scrambled regions 1, 2 or 3 of MIER1 and are shown in supplementary Fig. 1. The lane marked H2A/H2B contains *E. coli* expressed H2A:H2B complex purified on a Resource S column followed by Superdex S200 column.

Further confirmation of interaction was obtained using NMR spectroscopy. ^15^N-labelled MIER1(aa:1-177) was expressed in *E.coli* and titrated with unlabelled H2A:H2B dimer (expressed separately). The limited ^1^H chemical shift distribution of MIER1 in an HSQC spectrum is typical of an intrinsically disordered protein (Fig. 3b). Following addition of the H2A:H2B dimer, the HSQC spectrum clearly shows both chemical shift perturbations and a general broadening consistent with MIER1 binding the folded histone dimer (Fig. 3c). Approximately half the amino acids in MIER1(aa:1-177) appear to be involved in the interaction.

### Histone binding activity maps to two regions of conservation in the amino-termini of MIER proteins

To investigate which regions of the amino-terminus of MIER1 mediated the interaction with H2A:H2B, we hypothesised that regions conserved between MIER1, MIER2 and MIER3 would be most likely to mediate the interaction. Sequence alignment of the three isoforms reveals 3 regions with moderate conservation (Fig. 2d). Deletion of either region 1 or 2, but not 3, resulted in failure of MIER1 to co-purify with the H2A:H2B heterodimer (Fig.s 3d-g, Supplementary Fig. 1). Since regions 1 and 2 contain many negatively charged residues, it is important to rule out a non-specific interaction with the positively charged histone dimer. Therefore, to confirm that this interaction is specific we made MIER1 amino-terminal constructs in which the order of amino acids in regions 1 or 2 was scrambled. In both cases, MIER1 no longer co-purifies with the H2A:H2B dimer confirming a sequence-specific interaction (Fig. 3h-i, Supplementary Fig. 1).

To obtain a structural model of the interaction of MIER1(aa:17-75) regions 1 and 2 with H2A:H2B, we used Alphafold2 multimer to predict the mode of binding (Fig. 4a-c). The resulting model is structurally convincing and reveals how the MIER1 sequence has complementary stereochemistry to the H2A:H2B dimer surface. Overall the interface area between MIER1 and the histone dimer is 2,005Å^2^ (*23*). Region 1 is predicted with high confidence and adopts an extended configuration interacting with H2B through a conserved phenylalanine interacting in a non-polar pocket with residues Y43, I55 and Met60 in H2B. Region 2 is also predicted with high confidence, to form a helix making numerous non-polar interactions with the other end of H2B. Between regions 1 and 2 there are three additional helices containing many negatively charged residues that interact with the positively charged surface of the H2A:H2B dimer (Fig. 4f). This surface interacts with DNA in the assembled nucleosome (Fig. 4g).

**Fig. 4.**
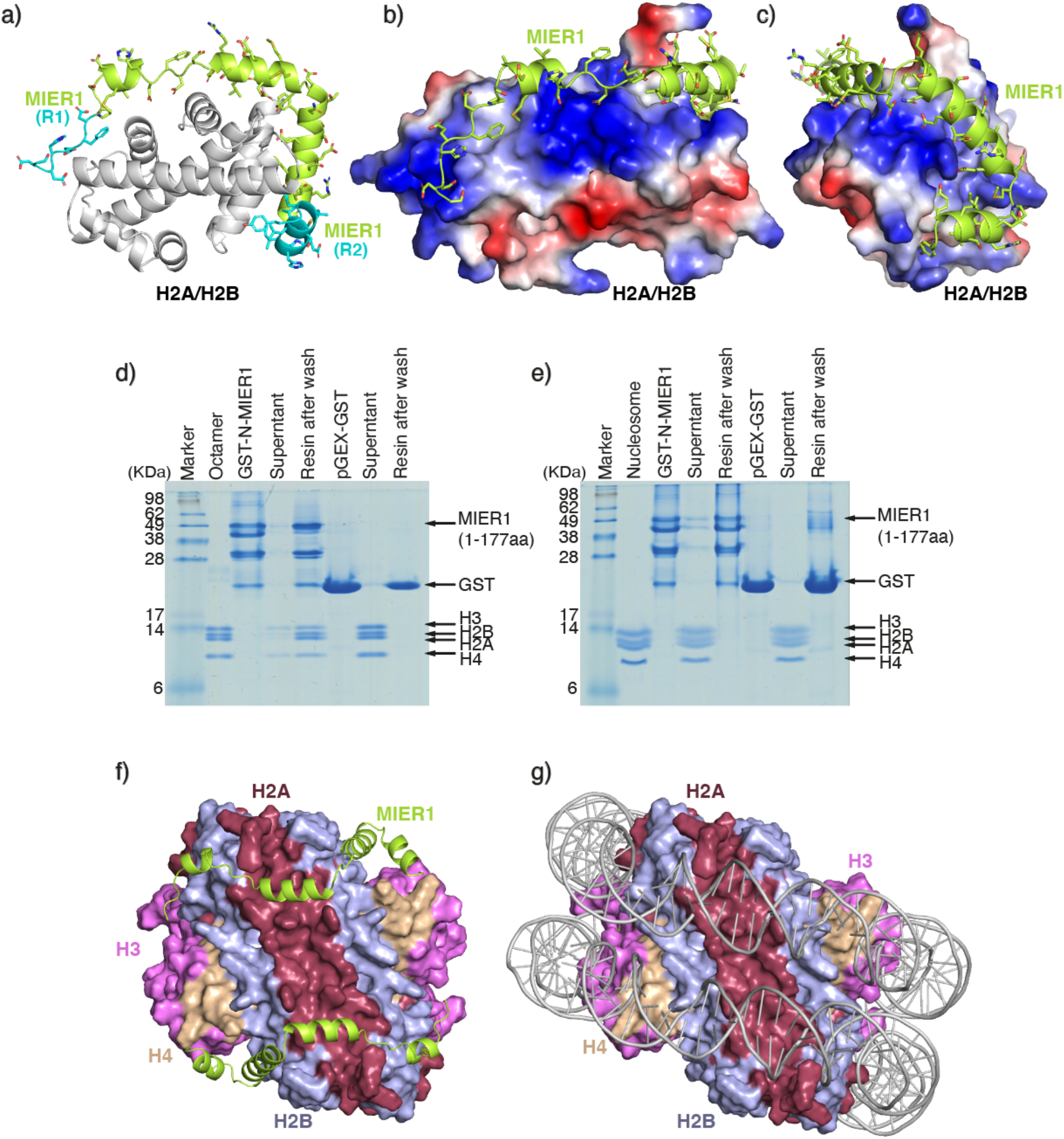
The amino-terminal region of MIER1(aa:1-177) binds a full histone octamer. a) Alphafold2 model of the interaction of MIER1(aa:17-75) (lime green) with an H2A:H2B dimer (grey cartoon). b) Alphafold2 model of the the interaction of MIER1(aa:17-75) (lime green) binding to H2A:H2B (electrostatic surface). c) Panel b rotated 260 clockwise about y-axis. d) GST pull down experiments confirm the ability of the amino-terminus of MIER1(aa:1-177) to bind a full histone octamer. e) GST pull down experiments show that the amino-terminus of MIER1(aa:1-177) cannot bind to a DNA-wrapped nucleosome. f) & g) Comparison of the binding of the amino-terminal region of MIER1(aa:17-75) to a histone octamer compared with a DNA-bound nucleosome.

### The MIER proteins can bind to H2A:H2B in the context of a histone octamer, but not when assembled into a nucleosome

As mentioned above, we noticed that the amino-terminal region of both MIER2(aa:1-178) and MIER2(aa:1-177) co-purified not only with H2A and H2B, but also with histones H3 and H4 (Fig. 2c). This suggests that the MIER proteins can bind to H2A:H2B in the context of a full histone octamer. Interestingly, the Alphafold2 model of MIER1 region1-2 (aa:17-75) bound to the H2A:H2B dimer appears to partially occlude the surface of H2B that interacts with histone H4 in the H3:H4 tetramer (Fig. 4c & f).

To explore this further, we expressed MIER1(aa:1-177) with a GST-tag in bacterial cells. This protein is significantly proteolyzed during purification from bacteria, consistent with being an intrinsically disordered protein. However, the degraded mixture is clearly able to pull-down a histone octamer, albeit with some dissociation of H3:H4, likely due to reduced stability of the octamer in low-salt buffer. In contrast, no histone octamer binding was observed for the GST-only control (Fig. 4d).

Given that we experimentally observed that the MIER1 amino-terminal region is able to bind to a histone octamer, we generated a further Alphafold2 multimer model including MIER1(aa:17-75) together with eight histone proteins. In this model MIER1 binds to the H2A:H2B dimer in the octamer context, exactly as the isolated H2A:H2B dimer, with the exception that the helix corresponding to the conserved region 2 is displaced and now interacts with histone H4. Fig. 4f shows the Alphafold2 model in the context of a full histone octamer compared with the DNA-wrapped nucleosome (Fig. 4g).

Importantly, neither GST-MIER1(aa:1-177), nor the GST-only control were able to interact with an intact DNA-wrapped nucleosome (Fig. 4e).

### The BAH domain from BAHD1 interacts with the MIER1:HDAC1 complex and recruits endogenous C1QBP

The Bromo-Adjacent-Homology-Domain containing protein 1 (BAHD1) has been identified as being a core component of the MIER1:HDAC1 complex (*13, 14*). BAHD1 consists of a carboxy-terminal BAH domain with an intrinsically disordered amino-terminal region contain a proline-rich region (Fig. 5a). We co-expressed the BAH domain from BAHD1(aa:525-780) together with the MIER1:HDAC1 complex confirming that the BAH domain is sufficient for interaction with the MIER1:HDAC1 complex (Fig. 5b).

**Fig. 5.**
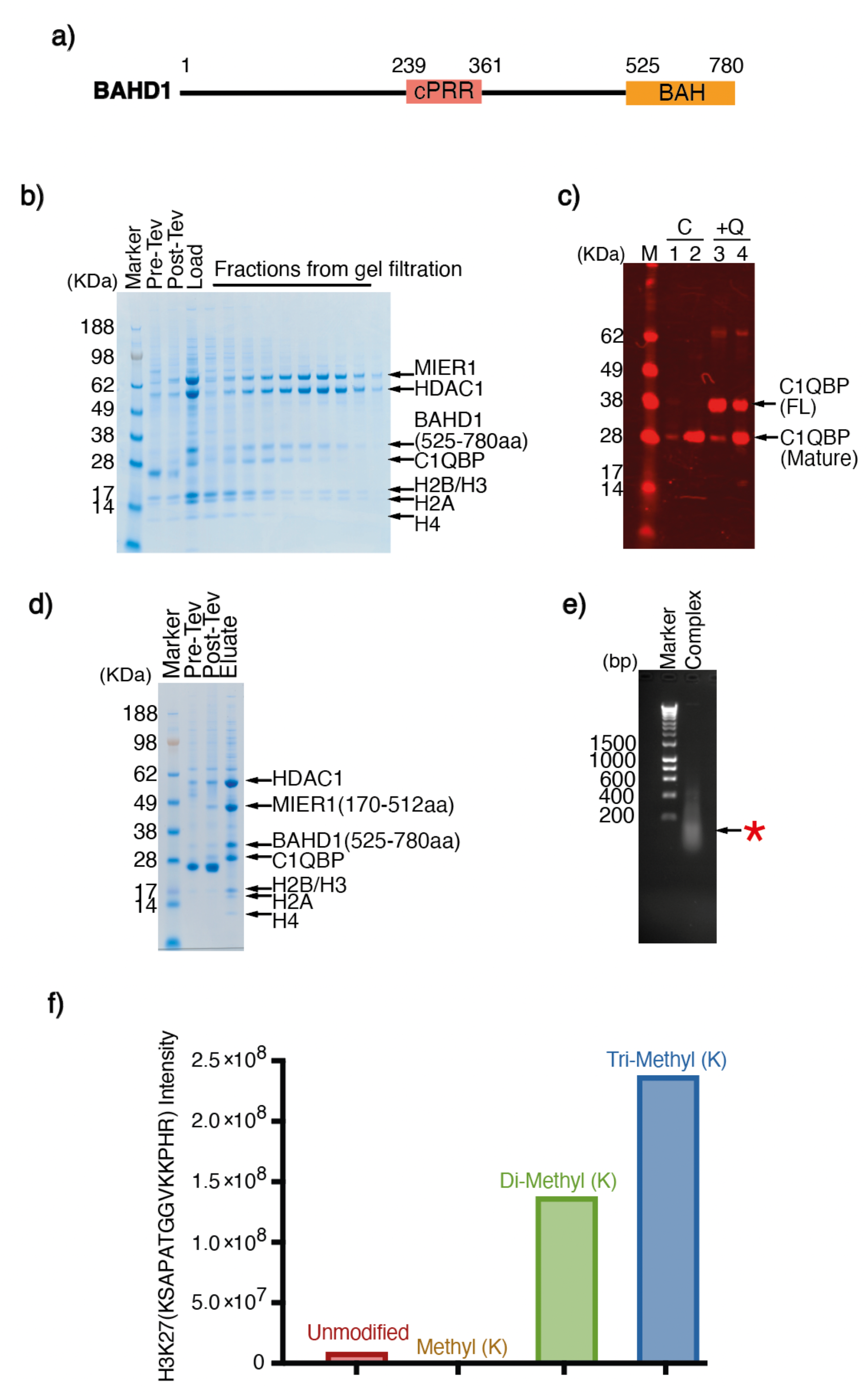
The MIER1:HDAC1:BAHD1 complex forms a stoichiometric complex with endogenous C1QBP and a nucleosome. a) Schematic of the domain structure of full-length BAHD1. b) The FLAG-tagged MIER1(aa:1-512):HDAC1: BAHD1(aa:525-780) complex was purified on anti-FLAG resin and a Superdex S200 column. The endogenous C1QBP and H2A:H2B:H3:H4 co-purified with the whole complex. c) Lanes 1 (lysate) and 2 (anti-FLAG purified) are from a co-transfection of MIER1(aa:171-512):HDAC1:BAHD1(aa:525-780) (labelled C). Lanes 3 (lysate) and 4 (anti-FLAG purified) are from a co-transfection of MIER1(aa:171-512):HDAC1:BAHD1(aa:525-780) with the addition of C1QBP (labelled +Q). The western blot was visualized using mouse anti-C1QBP antibody followed by goat anti-mouse 680RD. d) Co-purification of the MIER1(aa:171-512): HDAC1:BAHD1(aa:525-780) complex to show that the H2A:H2B:H3:H4 co-purified with MIER1 lacking the amino-terminus. e) Purified DNA obtained by micrococcal nuclease digestion of the MIER1(aa:171-512):HDAC1:BAHD1(aa:525-780):C1QBP:Nucleosome purified complex analysed in a 1xTBE 1% agarose gel. f) Quantitative MS analysis of histones co-purified with the MIER1(aa:171-512):HDAC1: BAHD1:C1QBP reveals that the majority of H3K27 is di or tri-methylated.

**Fig. 6.**
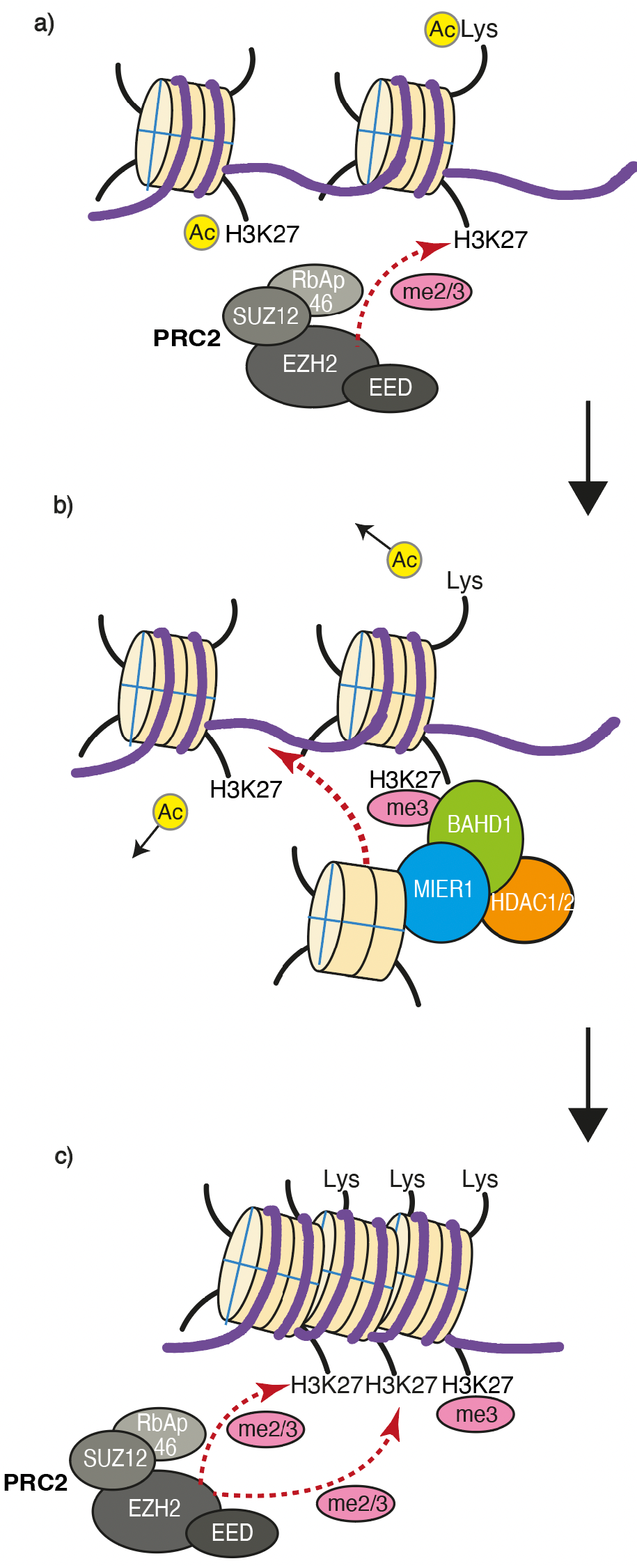
Proposed collaboration between the MIER1 and PRC2 complexes. a) PRC2 adds di-and tri-methyl marks to H3K27 to induce transcriptional repression. b) The MIER1 complex is recruited to chromatin through interaction with the H3K27me2/3 marks recognized by the BAH domain in BAHD1. c) This results in histone deacetylation and deposition of histone octamer feeding forward to enable the PRC2 complex to add further H3K27me2/3 marks.

Interestingly in these purifications we consistently observed an additional band with a molecular weight of around 28 kDa that co-purifies with a fraction of the complex. MS analysis of this gel band identified it as the protein C1QBP (aka: ASF/SF2-associated protein p32) (Supplementary Table 1). This 282aa protein is known to undergo a maturation process that removes a mitochondrial targeting sequence (aa:1-73) from the amino terminus leaving a mature protein of around 23kDa. To confirm the identity of this protein we performed a Western blot with an antibody against the full-length protein. Fig. 5c clearly shows that C1QBP is present in the lysate (lane 1) and strongly enriched in the purified complex (lane 2). To confirm the specificity of the antibody, we expressed a full-length C1QBP protein with an amino-terminal HA-tag that blocks maturation of the protein. This has an expected molecular weight of 33kDa (Fig. 5c, lane 3 and 4). From this it seems that the co-purified version of C1QBP is the mature form, consistent with the MS data (Supplementary Table 1) that did not identify any peptides from the amino-terminal 73 amino acids.

### The MIER1:HDAC1:BAHD1:C1QBP complex specifically recruits intact nucleosomes bearing H3K27me2/3 marks

Analysis of the gel filtration fractions following MIER1:HDAC1:BAHD1-BAH:C1QBP purification, shows that the earlier fractions contain all four histones H2A, H2B, H3 and H4 (Fig. 5b). In contrast later fractions contain only H2A:H2B. We initially hypothesised that a full histone octamer was binding to the amino-terminal region of MIER1 and that the presence of BAHD1:C1QBP might partly stabilise histone octamer binding so that it survives size exclusion chromatography.

To explore this further we co-expressed MIER1(aa:171-512) which lacks the amino-terminal histone octamer interaction region. We therefore expected that this complex would not recruit histone proteins. Unexpectedly, in this experiment we still co-purified all four histones (Fig. 5d). This suggests that the BAH domain from BAHD1 (or perhaps the co-purified C1QBP) is able to recruit either a histone octamer or a nucleosome. To distinguish these, we asked whether or not the histones that we co-purified in this complex are assembled with nucleic acid. We treated the peak fraction with micrococcal nuclease and analysed the released DNA on an agarose gel. The resulting band at around 150bp is consistent with nucleosomal DNA (Fig. 5e).

It has previously been shown that the BAH domain from BAHD1 recognises H3 histone tails bearing a K27me3 modification (*10*). We confirmed this interaction using a fluorescence anisotropy assay (Supplementary Fig. 2). We also used MS analysis to quantify the post-translational modifications on the histones that co-purified with the MIER1(aa:171-512):HDAC1:BAHD1-BAH:C1QBP complex (Supplementary Table 2). Importantly we found that the vast majority of histone H3 that co-purifies with the complex carries either di- and tri-methylation on H3K27, consistent with specific recognition by the BAH domain (Fig. 5f). Interestingly no acetylated lysine residues were detectable, consistent with efficient deacetylation by HDAC1 in the complex.

## Discussion

The MIER proteins recruit the deacetylase enzymes, HDAC1/2, via a conserved ELM2-SANT domain in the same way that the MIDEAS, MTA and RCOR proteins assemble the MiDAC, NuRD and CoREST complexes (*5-7*). Unlike the multimeric MiDAC and NuRD complexes, the MIER1 complex is monomeric (*5, 7*).

The finding that the MIER1 amino-terminal region has H2A:H2B / histone octamer binding activity suggests that the complex may play a role as a histone chaperone. This, in turn, suggests that the complex may be involved in either depositing or removing histones from chromatin. The Alphafold2 model of the MIER1 amino-terminal region bound to an H2A:H2B dimer makes excellent stereochemical sense, giving confidence that it is likely to be accurate. Importantly, the model also has numerous features in common with other peptide chaperones that bind H2A:H2B. Like ANP32E, Swr1, YL1, Chz1 and APLF, MIER1 has numerous acidic residues that interact with the DNA-binding surface of H2A:H2B (*24-28*). The MIER1 region (aa:17-75) overlaps closely with the other chaperones (except Chz1) which all have an aromatic residue (Y/F/W) that interacts with a non-polar pocket in the histone dimer (supplementary Fig. 3). MIER1(aa:17-75) follows a trajectory round the histone dimer similar to YL1 and Chz1 although the details of the interactions are different (*29*).

The experimental finding that MIER1 can bind to a complete histone octamer fits with the Alphafold2 model, but requires a reorientation of the region 2 alpha helix to avoid occluding the surface of the H2B that interacts with histone H4. This reorientation was seen in the Alphafold2 model of MIER1 bound to a complete octamer. Experimental structural biology would be required to confirm these models. The ability of a peptide to act as a chaperone for an intact histone octamer has been observed before for the APLF peptide (*28, 30*). The MIER1:octamer complex would appear to allow initial binding of DNA to the H3:H4 dyad position. We presume the DNA could then wrap around the octamer in both directions displacing the MIER1 complex.

The chaperone activity of MIER1 is unexpected and it appears that, of the six known HDAC1/2 complexes, only the MIER complex has this activity. Importantly, this activity is common to all three MIER proteins. Indeed, MIER homologues in D. melanogaster (Uniprot A1Z6Z7) and C. elegans (Uniprot P91437) show 47% and 37% identity compared with MIER1, respectively, in the amino-terminal region that we have shown interacts with the H2A:H2B histone dimer. This suggests that histone chaperone activity is conserved (Supplementary Fig. 4). Interestingly, MIER1 knockout mice are viable, albeit with metabolic dysfunction (*13*). It may of course be that MIER2 and MIER3 are able to compensate for the loss of MIER1 in these animals and that mice lacking all three genes would not be viable.

Genome-targeted HDAC complexes typically contain domains or sequence motifs, or accessory proteins that mediate interaction with chromatin (*31*). The MIER complex has been reported to associate with the chromodomain protein CDYL and the BAH domain protein BAHD1 (*10, 13, 32*). When we co-expressed MIER1 with HDAC1 and the BAH domain from BAHD1 we found that the endogenous protein C1QBP co-purified with the complex in apparently stoichiometric amounts. This association of C1QBP with the MIER1 complex is unexpected and raises questions as to its role. C1QBP is an enigmatic, multi-functional protein present in multiple cellular compartments (*33*). It has an amino-terminal 73 residue mitochondrial-targeting sequence that is cleaved during maturation (*34*). The carboxy-terminal 208 residues form a trimeric ring, doughnut-like structure (*35*). It has been reported to have roles in the regulation of apoptosis, pre-mRNA-splicing, mitochondrial protein synthesis and to act as a receptor for C1q at the cell membrane (*36, 37*). Perhaps more relevant for the MIER complex, C1QBP has also been reported to have a role in transcriptional regulation and homologous recombination (*38*). A recent report suggests that, intriguingly, C1QBP can act as a chaperone of histones H3 and H4 (*39*). This is supported by interactome analyses that identified interactions with all four cores histones and, interestingly, four proteins in the PRC2 repression complex (*40*).

Both this study and previous studies by others, suggest that the MIER1 complex is recruited to chromatin via interaction of the BAHD1-BAH domain with H3K27me3 marks (*10*). These marks are deposited by the PRC2 repressor complex leading to heterochromatin formation and gene repression. Interestingly, MIER1 has also been reported to form a complex with CDYL (*13, 32*), another reader of H3K27me2/3 marks. The finding that a deacetylase complex is recruited to this mark, suggests a feed-forward mechanism through which HDAC1/2 removes proximal H3K27ac marks, and perhaps other Kac marks such as H3K9ac (*41*), shutting down transcription and facilitating further methylation by PRC2. Since acetyllysine marks are associated with promoters and enhancers that typically have reduced nucleosome occupancy, it seems likely that the histone chaperone activity of the MIER complex allows it to deposit histone octamer at these sites thereby increasing nucleosome density and contributing to the formation of heterochromatin.

The intriguing observation of the juxtaposition of a deacetylase enzyme together with a histone octamer chaperone and H3K27me2/3 nucleosome-binding activity is both unexpected and unprecedented, but fits well with the role of the MIER complex in shutting down active transcription.

## Materials and Methods

### Mammalian protein expression and purification

MIER1 constructs were cloned into pcDNA3.0 expression vectors containing a amino-terminal 10xHis-3xFLAG tag followed by a TEV protease cleavage site. Full length HDAC1(aa:1–482) and BAHD1(aa:525-780) were cloned without affinity tags into the same vectors. Constructs we co-transfected in HEK293F suspension grown cells (Invitrogen) using polyethylenimine (PEI; Sigma) as a transfection reagent and harvested after 48 hours. Cells were lysed in buffer containing 50 mM Tris/HCl pH 7.5, 50 mM potassium acetate, 5% v/v glycerol, 0.4% v/v Triton X-100 and Complete EDTA-free protease inhibitor (Roche). The lysate was clarified by centrifugation and applied to Anti-FLAG M2 affinity resin (Sigma Aldrich) for 30 mins and washed three times with 50 mM Tris/HCl pH 7.5, 50 mM potassium acetate and 5% v/v glycerol and three times more with 50 mM Tris/HCl pH 7.5, 50 mM potassium acetate and 0.5 mM TCEP before being cleaved by TEV protease overnight. The protein complexes were purified further by gel filtration using a Superdex S200 column (Cytiva) with buffer containing 25 mM Tris/HCl pH 7.5, 50 mM potassium acetate and 0.5 mM TCEP.

### *E.coli* protein expression and purification

Truncated MIER1 was cloned into pGEX expression vectors containing a amino-terminal GST-tag followed by a TEV protease cleavage site and the recombinant protein was over expressed in *E. coli* strain Rosetta (DE3). Bacterial cultures were grown at 37°C to an optical density of ∼ 0.6, the temperature of the culture was reduced to 20°C and protein expression was induced by addition of isopropyl-β-d-thiogalactopyranoside (IPTG) to a final concentration of 40 µM. The cells were harvested after the cultures were grown for a further 16 hours. The cells were lysed in lysis buffer containing phosphate buffered saline (PBS), 1% v/v Triton X-100, 1 mM Dithiothreitol (DTT) and Complete EDTA-free protease inhibitor (Roche). The soluble fraction was then incubated with Glutathionine Sepharose (Cytiva) resin for 1 hour at 4°C with gentle agitation. The protein-bound resin was then washed three times with wash buffer (1× PBS, 1% v/v Triton X-100, 1 mM DTT) and then three times more with cleavage buffer (1× PBS, 1 mM DTT). The GST-tag was removed by incubation with TEV protease overnight at 4°C. The TEV cleaved complex was concentrated to around 500 μl using a 4 ml Amicon® Ultra centrifugal filter concentrator (Merck Millipore) with a membrane molecular weight cut off 10 kDa. The concentrated protein was filtered through a 0.22 μm centrifugal filter (Merck Millipore) prior to loading onto the gel filtration column. The protein sample was purified on a Superdex S200 (10/300) GL column (Cytiva) with buffer containing 25 mM Tris/Cl pH 7.5, 50 mM potassium acetate and 0.5 mM TCEP. 500 μl fractions were collected and 10 μl of the sample of the fractions was taken for analysis on NuPAGE 4%-12% Bis-Tris gel (Invitrogen). Fractions containing the purified protein were concentrated using a 0.5 ml Amicon Ultra centrifugal filter (Merck Millipore) with a membrane molecular weight cut off 10 kDa.

### Histone purification

Human histones cloned into pET3a (plasmids were a gift from Martin Browne and Andrew Flaus NUI Galway) were expressed separately in Rosetta2 (DE3) pLysS (Novagen). Cell pellets were resuspended in 30 ml histone wash buffer which contained 50 mM Tris/Cl pH 7.5, 100 mM NaCl, 5 mM DTT and Complete EDTA-free protease inhibitor (Roche). After sonication, the insoluble fractions were pelleted by spinning at 4 °C at 30,000 g for 15 mins and washed twice with wash buffer containing 50 mM Tris/Cl pH 7.5, 100 mM NaCl, 5 mM DTT, 1% Triton X-100 and Complete EDTA-free protease inhibitor (Roche) and three times with wash buffer without Triton X-100. To resuspend the inclusion bodies 0.5 ml of DMSO was then added and incubated at room temperature for 30 mins on a roller then 5 ml unfolding buffer containing 7 M Guanidine HCl, 20 mM Tris/Cl pH 7.5 and 10 mM DTT was added, and the incubation continued for an additional 1 hour.

After the pellet was fully dissolved, 70 ml 7 M Urea, 50 mM Tris/Cl pH 7.5 and 1 mM EDTA was added and the sample was centrifuged at 35,000 x g for 20 mins. The histone solution was then filtered through a 0.22 μm filter prior to loading onto a HiTrap Q HP. The flow through was then loaded onto a HiTrap SP HP. The histones were eluted with a gradient to 2 M NaCl. Fractions containing histone were then dialysed with three changes into buffer containing 0.1% acetic acid and 5 mM DTT. They were then freeze dried in aliquots.

### Histone octamer refolding and nucleosome reconstitution

H2A, H2B, H3 and H4 were dissolved in unfolding buffer which contained 7 M guanidine, 20 mM Tris pH 7.5 and 10 mM DTT and mixed with a small excess of H2A and H2B. The histone mix was dialysed against high salt buffer (20 mM Tris pH 7.5, 2.0 M NaCl, 1 mM EDTA and 5 mM mercaptoethanol) and then purified by size exclusion chromatography with a Superdex S200 column.

For the nucleosome reconstitution, 157 bp of 601 DNA was amplified by PCR from a 601 DNA template (*42*). The histone octamer and 157 bp DNA were mixed in equimolar amounts in 15 mM Hepes pH 7.5, 2 M KCl, 5% glycerol, 0.1% Triton X-100, 0.5 mM TCEP and dialysed stepwise to 15 mM Hepes pH 7.5, 50 mM KCl, 5% glycerol, 0.1% Triton X-100, 0.5 mM TCEP.

### Size exclusion chromatography with multi-angle light scattering (SEC-MALS)

Gel filtrated pure MIER1(aa:171-350):HDAC1 complex was reapplied to a Superdex S200 column and monitored with an Optilab T-rEX differential Refractive index detector connected to a DAWN HELEOS MALS detector (Wyatt Technology). The mass of the complex was calculated using ASTRA software version 6.1.

### Boc-Lys(Ac)-AMC HDAC Assay

A fluorescent based HDAC assay was used to determine HDAC activity of the MIER1 complex with BOC-lys(Ac)-AMC (BaChem) as the substrate. For these experiments 20 nM MIER1(aa:171-350):HDAC1 complex, with100 μM Ins(1,2,3,4,5,6)P_6_ or 30 μM SAHA or 30 μM MS275 in 25 mM Tris/Cl, pH 7.5 and 50 mM NaCl were incubated in the shaking incubator at 37°C for 30 mins before adding 100 μM substrate for a further 30 mins. The assay was developed by the addition of developer solution (2 mM TSA, 10 mg/ml Trypsin, 50 mM Tris pH 7.5, 100 mM NaCl). Fluorescence was measured at 335/460 nm using a Victor X5 plate reader (Perkin Elmer). Data was analysed using GraphPad Prism 9.0.

### Isothermal titration calorimetry (ITC)

The ITC experiment was performed using a VP-ITC instrument (MicroCal). H2A/H2B dimer (4.6 μM) was loaded in the sample cell and 170 μM of MIER1(aa:1-177) in the syringe. The titration experiment was performed at 25°C and consisted of 30 injections of 5 μl each with a 5 min interval between additions. The raw data were integrated, corrected for nonspecific heat, normalized for the molar concentration and analysed according to a 1:1 binding model.

### NMR measurement

For NMR sample preparation, MIER1(aa:1-177) was uniformly labelled by overexpression in M9 minimum medium, containing ^15^NH_4_Cl as the sole nitrogen source. The expression and purification of labelled MIER1 was performed as described above. MIER1(aa:1-177) (200 μM) with 10% D2O in a final volume of 330 μl of PBS buffer was placed in 3 mm NMR tubes (Norell). (^15^N-^1^H)-HSQC spectra were measured on a Bruker AVIII-600 MHz spectrometer equipped with a cryoprobe. Data were processed using Topspin (Bruker) and analysed with Sparky. H2A:H2B dimer was added to the labelled MIER1(aa:1-177) protein at the ratio of 1:1 and incubated for 30 mins at 4°C. The MIER1(aa:1-177):H2A:H2B complex was then purified on a Superdex S200 column before collecting (^15^N-^1^H) HSQC NMR data.

### Western Blot

Western blotting was carried out using a NuPAGE 4-12% Bis-Tris gels (Invitrogen) and a semi-dry transfer system using nitrocellulose membranes. Membranes were blocked for 1 hour at room temperature using 5% skimmed milk TBS/T and incubated with the primary mouse antibody C1QBP (1:1000 H-9:sc-271200, Santa Cruz) diluted in blocking buffer overnight at 4°C. Secondary antibody goat anti-mouse 680RD (1:10,000, LI-COR), was incubated with membranes for 1 h at room temperature and detected using an Odyssey CLx digital imaging system (LI-COR).

### GST-Pull down experiment

GST-MIER1(aa:1-177) and GST tag alone were purified separately without eluting the proteins from the Glutathione Sepharose resin (Cytiva) in PBS, 1mM TCEP and Complete EDTA-free protease inhibitor (Roche). Excess octamer and nucleosome were mixed separately with the GST-tagged MIER1 or with the GST-tag alone and incubated for 30 mins at 4°C. After washing 5 times with PBS, 0.5 mM TCEP, the samples were analysed on an 18% SDS-PAGE.

### DNA extraction

50U/μl micrococcal nuclease and 10 mM CaCl_2_ was added to the purified MIER1(aa:171-512):HDAC1:BAHD1(aa:525-780):C1QBP complex. This was incubated for 30 mins at 37°C and EDTA to 10 mM was added to stop the reaction. 10 µl of proteinase K with 1% SDS were added and incubated for 30 mins at 37°C. This was then adjusted to 1M NaCl and Phenol:Chloroform:Isoamyl Alcohol extracted. The DNA was ethanol precipitated, analysed on a 1x TBE, 1% agarose gel and visualised with ethidium bromide.

### Fluorescence polarisation assay

Fluorescence polarisation experiments were performed using fluorescently 5-FAM labelled histone peptides (Cambridge Research Biochemicals).

H3 1-19 ARTKQTARKSTGGKAPRKQ-ed-[4-Abu]-[5-FAM]

H3 1-19 K4me3 ART-[Lys(Me3)]-QTARKSTGGKAPRKQ-ed-[4-Abu]-[5-FAM]

H3 1-19 K9me3 ARTKQTAR-[Lys(Me3)]-STGGKAPRKQ-ed-[4-Abu]-[5-FAM]

H3 16-35 PRKQLATKAARKSAPATGG-ed-[4-Abu]-[5-FAM]

H3 16-35 R26me2a PRKQLATKAA-[ADMA]-KSAPATGG-ed-[4-Abu]-[5-FAM]

H3 16-35 K27me3 PRKQLATKAAR-[Lys(Me3)]-SAPATGG-ed-[4-Abu]-[5-FAM]

(ADMA: asymmetric dimethyl arginine; Abu: aminobutyric acid; ed: ethylene diamine linker)

The assays were performed in black non-binding surface 384-well assay plates (Corning Life Sciences). Multiple titrations were performed using a fixed concentration of 5 nM histone peptide with increasing concentrations of *E. coli* expressed BAHD1(aa:588-780) (0-142µM) in a final volume of 25µl in 25 mM Tris/Cl, pH 7.5 and 50 mM NaCl. The plate was incubated in the shaking incubator for 15 mins at 37°C, then centrifuged at 61 g for 10 seconds and the data were acquired in a Victor X 5 plate reader (Perkin Elmer) with an excitation wavelength of 480 nm and an emission wavelength of 535 nm. The subsequent data were analysed with GraphPad Prism 9.0. Kd values were calculated by nonlinear curve fitting with a one-site binding model.

### Mass spectrometry analysis

#### Propionylation of histones and in-gel digestion

Gel bands were derivatised according to (*43, 44*) with some modifications. Briefly, 50 µl of 50 mM ammonium bicarbonate was added to the gel pieces and incubated with 16.6 µl of propionylation reagent (1:3 v/v propionic anhydride in acetonitrile) for 15 mins at 37°C with agitation in a thermomixer at 900 rpm (Eppendorf, UK). The supernatant was then removed and the derivatisation process was repeated. After discarding the supernatant, the gel was destained, and proteins were reduced with 10 mM dithiothreitol for 1 hour at 55°C and then alkylated with 55 mM iodoacetamide for 20 mins at room temperature in the dark. Protein in the gel was digested with 100 µl of trypsin (0.02 µg/µl) at 37°C overnight. Peptides were then extracted twice with 100 µl of 3.5% formic acid in 30% ACN then followed by 5% formic acid in 50% ACN. Eluted peptides were dried in a vacuum concentrator and stored at - 20 °C until MS analysis.

#### MS and data analyses

Peptides were resuspended in 0.5% formic acid and analysed using nanoflow LC-MS/MS using an Orbitrap Elite (Thermo Fisher) hybrid mass spectrometer equipped with a nanospray source, coupled to an Ultimate RSLCnano LC System (Dionex). Peptides were desalted online using a nano trap column, 75 μm I.D.X 20mm (Thermo Fisher) and then separated using a 120-min gradient from 5 to 35% buffer B (0.5% formic acid in 80% acetonitrile) on an EASY-Spray column, 50 cm × 50 μm ID, PepMap C18, 2 μm particles, 100 Å pore size (Thermo Fisher). The Orbitrap Elite was operated with a cycle of one MS (in the Orbitrap) acquired at a resolution of 120,000 at m/z 400, with the top 20 most abundant multiply charged (2+ and higher) ions in a given chromatographic window subjected to MS/MS fragmentation in the linear ion trap. An FTMS target value of 1e6 and an ion trap MSn target value of 1e4 were used with the lock mass (445.120025) enabled. Maximum FTMS scan accumulation time of 200 ms and maximum ion trap MSn scan accumulation time of 50 ms were used. Dynamic exclusion was enabled with a repeat duration of 45 s with an exclusion list of 500 and an exclusion duration of 30 s. Raw mass spectrometry data were analysed with MaxQuant version 1.6.10 (*45*). Data were searched against a human UniProt reference proteome (downloaded June 2019) using the following search parameters for standard protein identification: enzyme set to Trypsin/P (2 mis-cleavages), methionine oxidation and N-terminal protein acetylation as variable modifications, cysteine carbamidomethylation as a fixed modification. For histone PTM analysis the following settings were used: enzyme set to Trypsin/P with up to 5 missed cleavages, methionine oxidation, lysine propionylation, lysine acetylation, lysine mono, di and tri-methylation were set as variable modifications, cysteine carbamidomethylation as a fixed modification. A protein FDR of 0.01 and a peptide FDR of 0.01 were used for identification level cut-offs based on a decoy database searching strategy.

## Acknowledgments

We are grateful to the PROTEX facility at the University of Leicester for preparation of expression constructs.

## Author contributions

Conceptualization: SW, JWRS

Methodology: SW, LF, KP, TJR, DV, MC, CD

Investigation: SW, LF, KP, TJR, DV, MC, CD

Supervision: JWRS, MC, CD

Writing—original draft: SW, JWRS

Writing—review & editing: SW, LF, MC, CD, JWRS

**Fig. S1.**
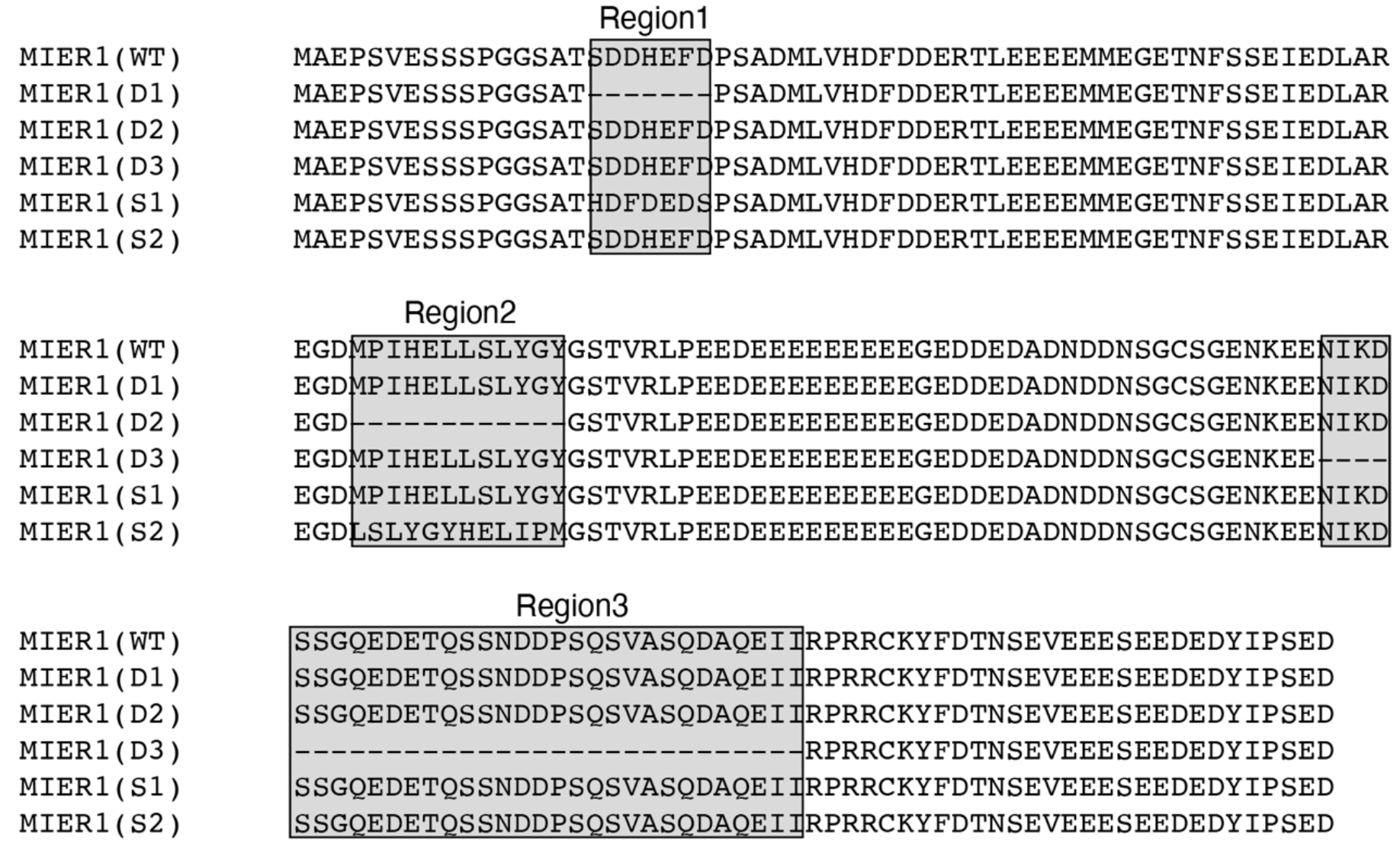
Sequences of WT MIER1 (1-177), sequences of the deletions of Regions 1, 2 and 3 (D1, D2 and D3) and sequences of the scrambled regions 1 and 2 (S1 and S2).

**Fig. S2.**
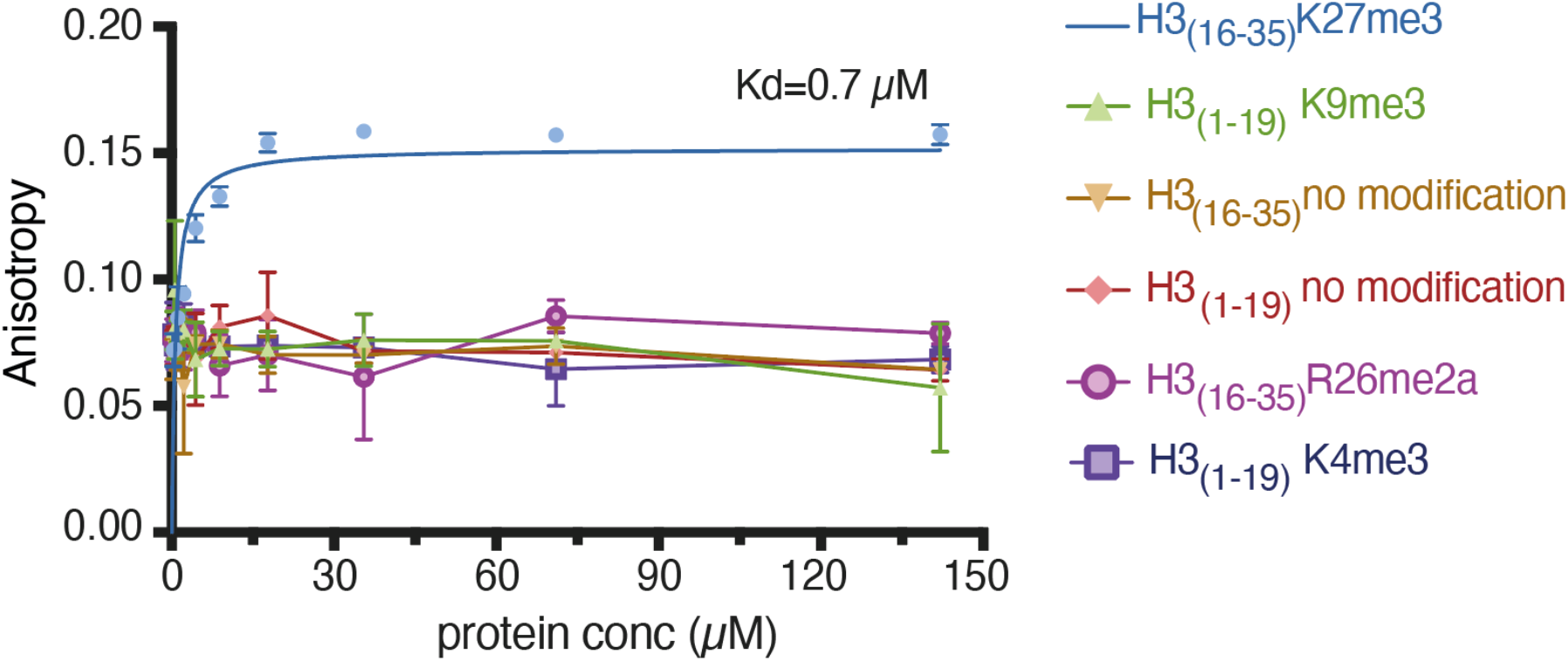
Fluorescence anisotropy assay of the binding of the various modified histone peptides to BAH domain of BAHD1. Error bars indicate ± s.e.m (n=3).

**Fig. S3.**
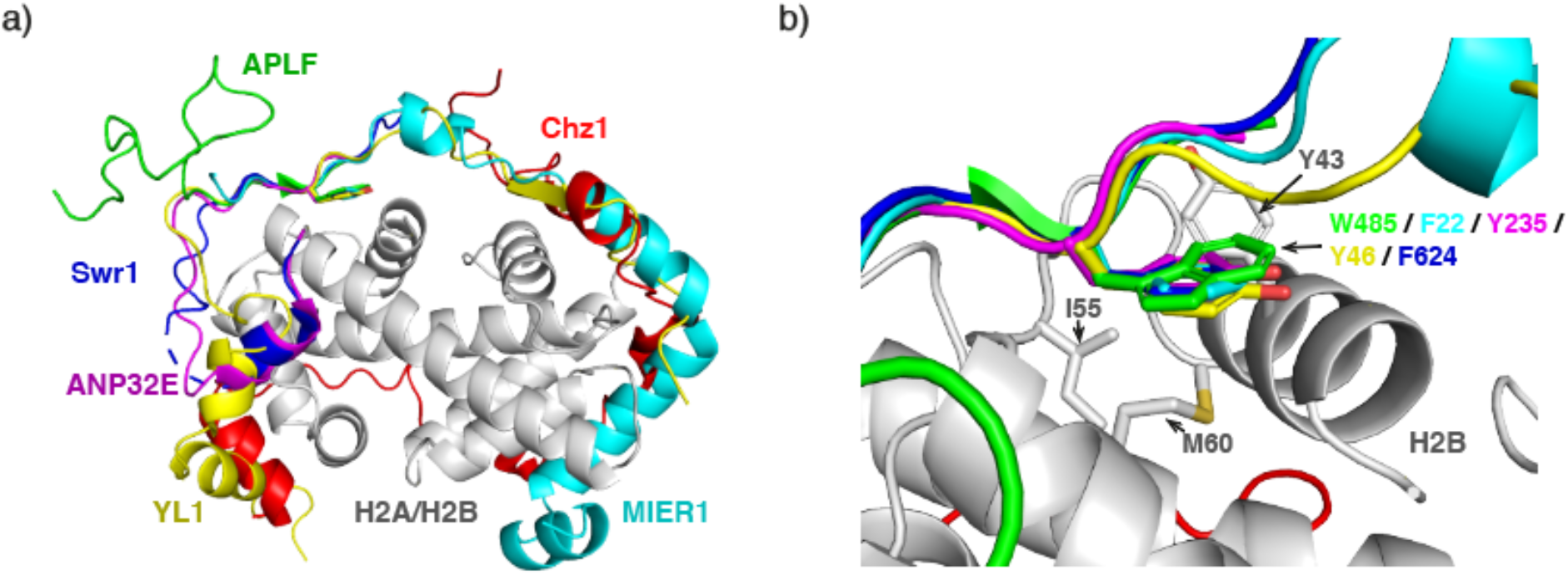
a) Comparison of the Alphafold2 multimer model of MIER1(aa:17-75), with Chz1 (PDB: 2JSS), Swr1 (PDB: 4M6B), ANP32E (PDB: 4CAY), YL1 (PDB:5FUG), APLF (PDB: 6YN1) all bound to a histone H2A:H2B dimer. b) Detail of the shared non-polar interaction with the H2A:H2B dimer – colors according to panel a.

**Fig. S4.**
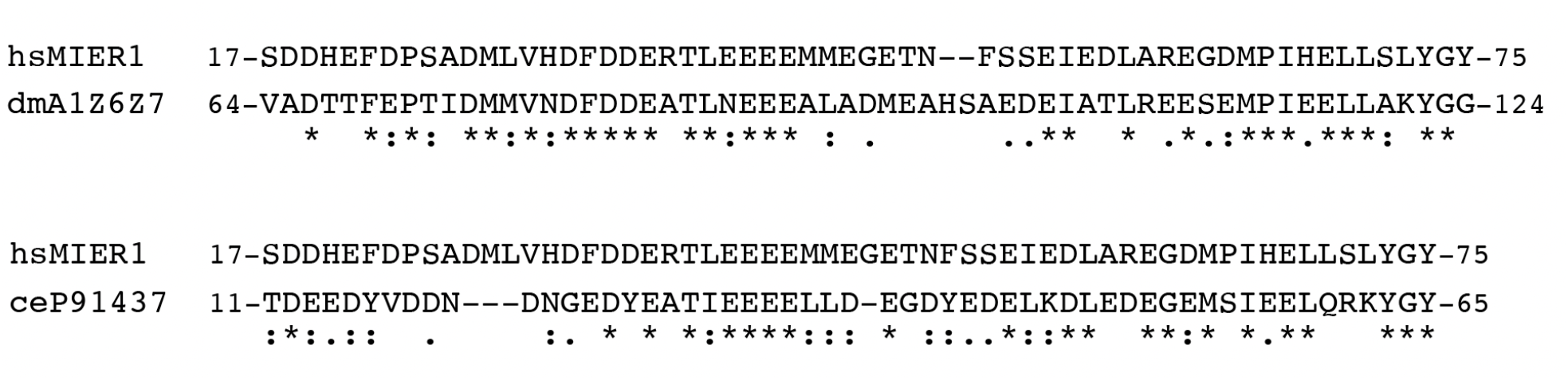
Alignments of the N-terminus of H. sapiens MIER1 (Uniprot Q8N108) with D. melanogaster MIER1 (Uniprot A1Z6Z7) and C. elegans MIER1 (Uniprot P91437)

**Table S1**.

Data related to Figure 5b. MS analysis of the MIER1:HDAC1:BAHD1 complex identifies C1QBP as a co-purifying protein.

Protein level data is contained in the “Protein groups” tab with complex components highlighted in yellow. Samples A-D correspond to gel bands containing HDAC1, MIER1, BAHD1 and C1QBP, respectively. Peptide data for C1QBP is included in the “C1QBP peptides” tab, with sequence coverage highlighted in red. All data is filtered to a 1% FDR at the protein and peptide level.

**Table S2**.

Data related to Figure 5f. Quantitative MS analysis of histones co-purified with the MIER1(171-512):HDAC1:BAHD1:C1QBP reveals that the majority of H3K27 is di or tri-methylated.

Protein level data is contained in the “Protein groups” tab. Identified acetyl, methyl, dimethyl and trimethyl lysine site identifications are listed in the relevant tabs. A summary of quantified levels of peptide (KSAPATGGVKKPHR) modification forms encompassing H3K27 is shown in the “H3K27 modification state” tab. All data is filtered to a 1% FDR at the protein and peptide level.

